# Robust parameterization of a viral-immune kinetics model for sequential Dengue virus (DENV) infections with Antibody-Dependent Enhancement (ADE)

**DOI:** 10.1101/2023.08.29.555313

**Authors:** Joshua Macdonald, Hayriye Gulbudak

**Author notes:** Corresponding author: HG. Faculty of Life Sciences & School of Zoology, Tel Aviv University, Tel Aviv-Yafo, Israel (Joshua Macdonald).

## Abstract

Dengue (DENV), a neglected tropical disease, is a globally distributed arboviral (genus *Flavivirus*) pathogen primarily spread by *Aedes* mosquitoes and infecting approximately 390 million individuals annually. A challenge to successful control of DENV is that after primary infection (or vaccination) due to waning, secondarily infected patients (or vaccinated individuals) can have an elevated risk of severe Dengue due to a phenomenon known as antibody-dependent enhancement (ADE), that is: preexisting cross-reactive IgG antibody concentrations can increase dengue severity. In this study, we first robustly parameterize a unified within-host viral and immune kinetics model to viral kinetics data for serotypes DENV1, 2, and 3 collected at the Hospital for Tropical Diseases (Ho Chi Minh City, Vietnam) while allowing independent variation in infection start time among hosts. Our model recapitulates the data well, including cross-reactive antibody concentration-dependent enhanced severity in secondary infections, and captures empirically observed differences between primary and secondary DENV infections, such as time to peak viral load, duration of viremia, and maximum viral titer. Our parameterization also captures meaningful differences in serotype-specific kinetic parameters that drive these differences. Subsequently, we (i) show that variation in initial IgG antibody concentration is sufficient to mechanistically explain the observed differences between primary and secondary infection in terms of the time course of events across serotypes and (ii) leverage our modeling results paired with long-term NS1-specific IgG antibody decay data from Recife, Northeast Brazil, to estimate the half-life of Dengue IgG antibodies and the time frame of the risk window for escalated disease severity due to ADE.

## 2 Introduction

Dengue, a vector born RNA *Flavivirus* (DENV), is primarily spread by *Ades* mosquitoes and is thought to have originated approximately 1000 years ago in either Africa or Asia in monkeys and to have established endemic transmission in humans a few hundred years ago, with each serotype independently making the jump from its wild reservoir(s) to humans [1]. Though DENV is globally distributed, most cases occur in South America and Southeast Asia. Primary DENV Infection does not confer lifelong immunity due to the circulation of four antigenically distinct serotypes (DENV1, 2,3,4) [2, 3]. As such, in countries where DENV is endemic, many individuals will experience infections of multiple DENV serotypes in their lifetimes, and subsequent infections may be more severe. In this study, we turn our attention to three tasks: (i) robust serotype-specific parametrization of our previously developed viral and immune kinetics model [4] to differentiate life history characteristics of each serotype, (ii) demonstration of our model’s ability to capture empirically observed differences in the time course of events between primary and secondary infection, given this parameterization, *solely as a consequence of differences in concentration-dependent activation and replication rates*, and (iii) characterization of the “intermediate risk-window” of cross-reactive IgG antibody concentration [2] via estimation of concentrations associated with this risk, of the half-life of DENV IgG antibodies, and finally of the time-frame associated with this risk-window of concentrations.

Several empirically observed differences exist in the time course of events between primary and secondary infection. Invasion of the host via mosquito bite is thought to occur 4-6 *days* before symptom onset in both primary and secondary DENV patients; viremia to begin approximately 2 *days* before symptom onset and end around symptom day seven for primary infections and between 5-7 days after onset of symptoms for secondarily infected patients [3]. Peak viral load tends to occur more rapidly and at a higher concentration level in secondary DENV patients than in primary. However, on average, secondarily infected patients are also thought to have a relatively shorter period of viral proliferation (viremia) [3].

Clinical signs of DENV are diverse, including fever, shock, and hemorrhage [3]. Individual cases can range from asymptomatic (or unreported) [*≈*67% of infected individuals] to what is known as Dengue Hemorrhagic Fever (DHF) [*<* 5% of symptomatic individuals], characterized by pleural effusion due to vascular permeability [3]. After primary infection, immune memory is cross-protective for two to three months, after which protection is serotype specific [5], and secondarily infected patients have elevated risk of severe dengue [6]. No efficient DENV-specific antivirals exist, so developing effective vaccines is crucial to curtailing incidence rates and disease progression [7].

Uniquely among globally distributed pathogens, successful DENV vaccine development and deployment are challenged by a phenomenon known as Antibody-dependent enhancement (ADE), where differences in initial IgG antibody levels in secondarily infected (or vaccinated) hosts can enhance infection severity [2, 6–9]. These IgG antibodies may be broadly grouped into two classes: specific and non-specific (sero-cross-reactive), characterized by differences in targeted viral proteins (epitopes) within the capsid encapsulated viral package (virion), with the latter targeting epitopes common among DENV serotypes, and the former serotype-specific proteins [3].

ADE has been empirically observed in multiple cohorts of patients at different times and in different countries [2, 6, 8, 9]. There are multiple proposed mechanisms for ADE. Neutralization of virions requires higher concentrations of cross-reactive antibodies relative to the specific response [10]. Thus, the “multiple-hit” stoichiometry requirement hypothesis [11–13] suggests that a threshold number of cross-reactive antibodies bound to a virion are required for neutralization. Antibody-virion binding can also increase the probability of cell infection [10], thereby effectively enhancing the viral replication rate. Another possible mechanism of ADE is “original antigenic sin,” where antibody populations compete and interfere relatively more with efficient IgG activation in secondary infection [14]. In addition to the increased risk of severe Dengue for secondarily infected DENV patients, differences in initial IgG antibody concentration and the resulting concentration and time-dependent net viral replication and IgG activation rates may also be sufficient to mechanistically explain observed differences between primary and secondary infection in the timing of within-host events [3].

Mechanistic models encapsulating various aspects of these possible mechanisms have been used to study Dengue infection at either the within-host and between-host scales [15–18]. These efforts include capturing multi-strain epidemiological dynamics with potential secondary infection and ADE due to partial or temporary cross-reactivity [19–22]. In all of these population-scale models, ADE has been incorporated through parameters associated with secondary infection. *However, the empirical evidence points to constitutive cross-reactive IgG antibody levels at the start of secondary infection as the determinant of infection severity* [2, 3, 6, 8, 9]. Similarly, at the within-host scale, several models have incorporated ADE effects [17, 23, 24], *but these models failed to produce enhanced severity dependent on initial cross-reactive IgG concentration* [4]. Therefore, initially inspired by [2], in [4], as part of a multi-scale modeling framework that links within-host viral and immune dynamics to epidemiological spread via a vector–host partial differential equation model structured by host antibody level, we formulated a viral-immune kinetics model which mechanistically recapitulates aspects of antibody-dependent enhancement in Dengue infection [2, 3]. In this work, we showed via simulation studies that the model captures several potential explanatory mechanisms for ADE [4], among them that *ADE occurs during an intermediate risk window for the decay of cross-reactive antibody titer*.

To our best knowledge, this is the first study to parameterize a unified model encapsulating ADE effects for sequential DENV infections among multiple serotypes of DENV and the first to characterize the intermediate risk window. In section 3, we develop a model parameterization approach that allows for a large degree of uncertainty in the early course of infection. Such an approach is necessary given the observational nature of the data, collected only after symptoms’ onset. We then use this approach to identify kinetic parameters for DENV1, 2, and 3 and identify differences in life history among serotypes. We then use these results paired with simulation studies to test our model’s ability to recapitulate the data and estimate resulting quantities to characterize these life histories, such as time to peak viral load and duration of viremia (section 4.1). Next, we demonstrate that differences in initial cross-reactive IgG antibody levels between primary and secondary infection across serotypes are sufficient to characterize empirically observed differences between primary and secondary infection (section 4.2). Finally, we characterize the intermediate risk window for enhanced severity secondary infection in terms of both cross-reactive IgG antibody concentrations and time spent in this risk window of concentrations (section 4.3).

## 3 Materials and methods

### 3.1 The model

In [4], we presented a simplified within-host viral and immune kinetics model that still captures the essential mechanisms of ADE. In this work, we considered IgG antibodies as proxies for the total immune populations, including innate (non-specific) and T-cell immune effectors, to model the host immune response. Let *x*(*τ*) denote virus concentration, *y*(*τ*) cross-reactive antibody concentration, and *z*(*τ*) specific antibody concentration, where *τ* is the time since infection began. In this model, we assumed that the virus undergoes exponential growth, parameter *r*, and the specific IgG response *z*(*τ*) clears the virus and saturates according to Michaelis-Menten kinetics. We included multiple mechanisms for ADE in this model. First, we captured differences in neutralization efficiency by modeling neutralization by cross-reactive response *y*(*τ*) with a sigmoidal Hill equation of (*n* = 2), representing positive cooperativity [25]. Thus encapsulating the “multiple-hit” stoichiometry requirement hypothesis [11–13]. We also included a Michaelis-Menten replication enhancement term dependent on cross-reactive antibodies *y*(*τ*). The “original antigenic sin,” mechanism where antibody populations compete and interfere with each other [14] was incorporated via interference competition coefficients *k*_1_ and *k*_2_ in the denominators of the hill function for *y* and *z* respectively. With these features in mind, this within-host model is novel for its enzyme kinetic mechanisms of ADE [4].

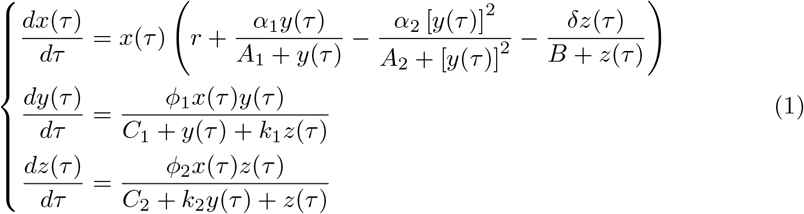

Here, for a specific serotype, we assume that, on average, the only difference between a primary and secondary infection in distinct patients in terms of kinetic parameters should be the difference in IgG antibody levels between the primarily and secondarily infected hosts at the start of an infection and the resulting concentration and time-dependent net viral replication and IgG activation, and clearance rates (see Fig. 4g-i):

**Fig 1.**
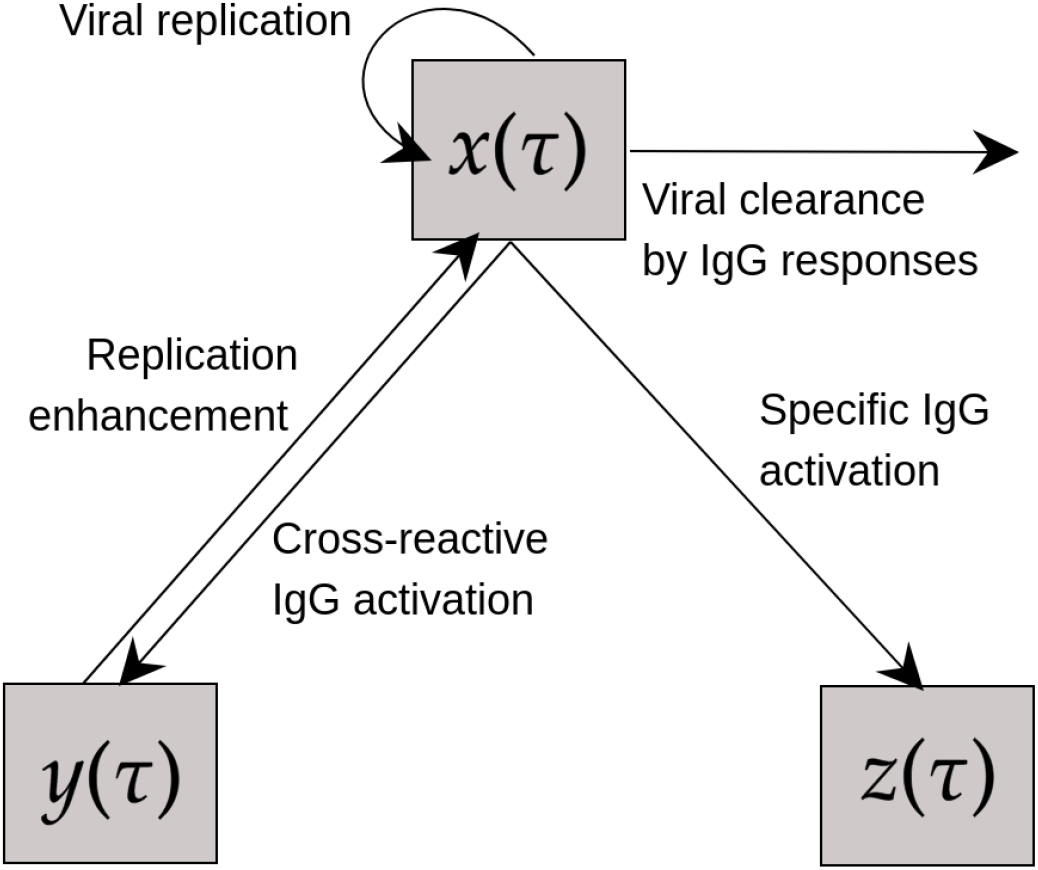
Model diagram. *x* is pathogen, *y* is cross-reactive IgG antibody, *z* is specific IgG antibody.

**Fig 2.**
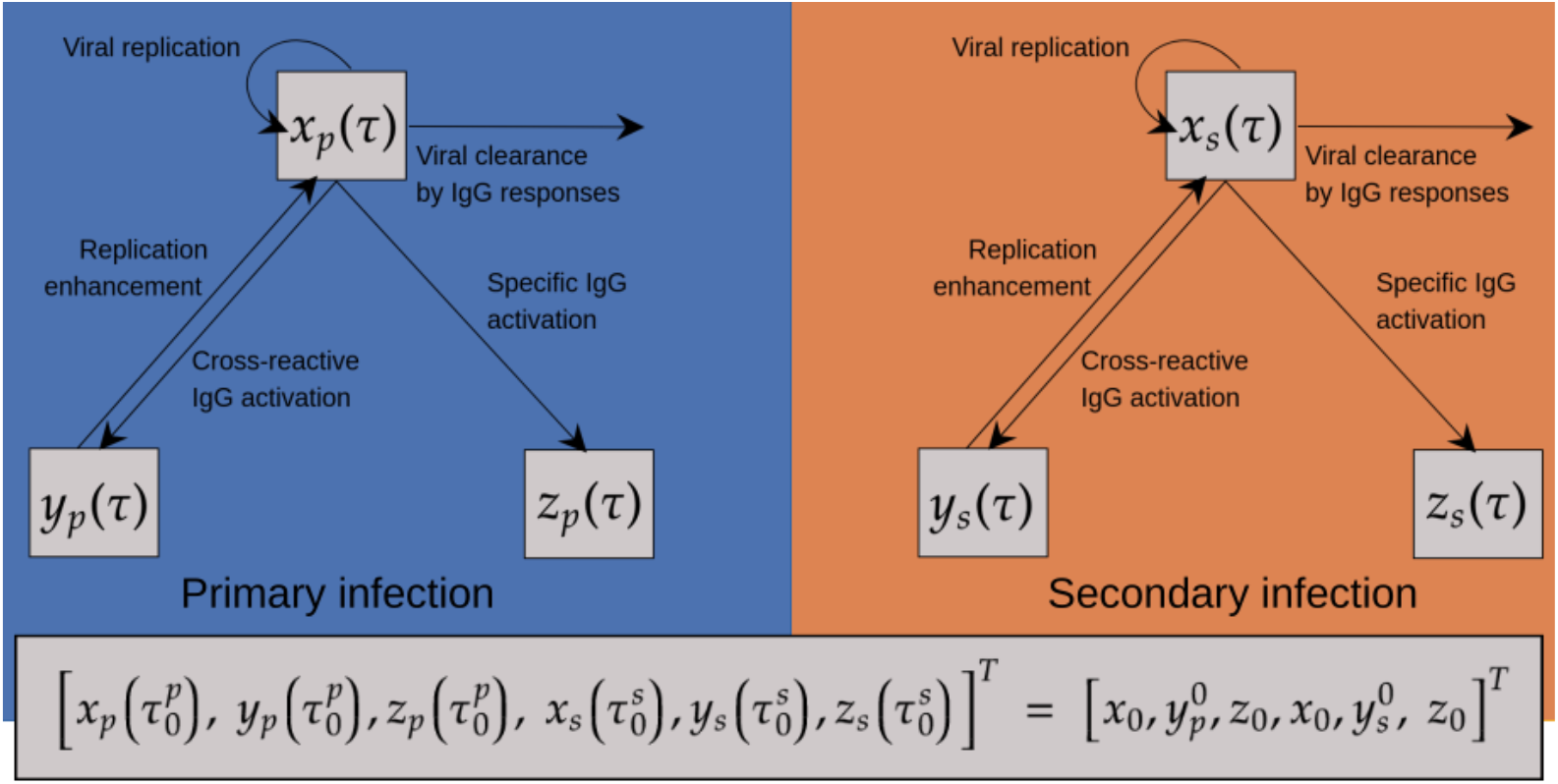
Diagram for the model (3). *x* is pathogen, *y* is cross-reactive IgG antibody, *z* is specific IgG antibody. The subscripts *p, s* denote primary and secondary infection, respectively.

Where *r*_*n*_, *y*_*A*_, *z*_*A*_ denote the concentration-dependent net viral replication, cross-reactive IgG activation, and specific IgG activation rates, respectively:

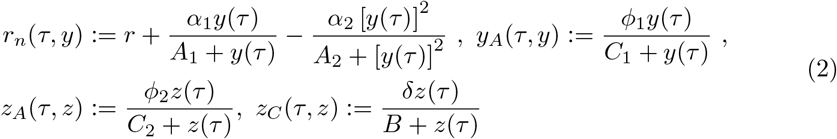

This results in the model:

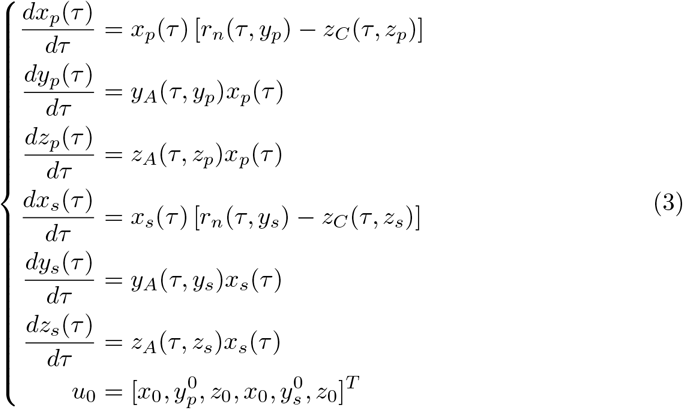

where subscripts *p, s* denote primary and secondary infection, respectively. 124

### 3.2 The data

#### Viral kinetics data

The viral load time series data were collected from the Hospital for Tropical Diseases (Ho Chi Minh City, Vietnam) between May 2007 and July 2008 as part of a clinical trial to determine if CQ inhibited viral replication of Dengue. Patients with illness histories of 72 hours or less were assigned randomly to either a 3-day course of CQ (*n* = 153) or placebo (*n* = 154). DENV infection was confirmed in 257 (84%) of the patients. The primary endpoints were time to resolution of DENV viremia and time to resolution of DENV NS1 antigenemia [27]. No statistically significant effect of CQ was detected on the duration of viremia [27], and the data from both treatments have been analyzed together previously [16, 27], an approach which we also follow. Among confirmed positive patients, Dengue viremia in plasma was measured using an internally controlled, serotype-specific, real-time RT-PCR TaqMan assay (see [28]), and results 136 were expressed as cDNA equivalents per ml of plasma [27]. Both primary (DENV1, *n* = 18; DENV2, *n* = 6; DENV3, *n* = 6; DENV4, *n* = 0) and secondary (DENV1, *n* = 124; DENV2, *n* = 45; DENV3, *n* = 33; DENV4, *n* = 7) patients took part in this trial, resulting in 2232, 270, and 198 combinations of patients res. Due to the lack of primary DENV4 data, we restrict ourselves to working with the DENV1, 2, and 3 data.

#### Long-term immune kinetics data

The long-term immune kinetics data [29] used to estimate the time frame of the intermediate risk window were gathered in Recife, Northeast Brazil, as part of a longitudinal cohort of 266 confirmed DENV3 positive individuals lasting more than three years post-infection. The Anti-dengue NS1-specific IgG antibody data were characterized by enzyme-linked immunosorbent assay (ELISA) [30]. Specifically, we considered individuals determined to have primary DENV infections [31] and had at least one data point past the maximum recorded antibody titer (*n* = 154).

### 3.3 Model fitting

From a data-driven modeling perspective, there are several challenges in parameterizing a mechanistic model incorporating primary and secondary DENV infections in a unified framework. First, early infection time data is not available for human hosts, both because such low viral concentrations will not be detectable by assays (cf. [16, 27]) and because, for obvious reasons, limitation to observational data, which typically begin within the first couple of days post symptom onset in individual hosts (cf. [27]), meaning that peak viral load for severe secondary infection is frequently not observed [3]. Additionally, given the observational nature of available data and the stochasticity of secondary infection, primary and secondary *in vivo* patient data is not generally available from the same hosts [27]. Finally, immune and viral kinetic data are often collected separately from patients for various reasons.

Over *M* = *N*_*p*_ *·N*_*s*_ iterations draw primary host *i* and secondary host *j*, 1 *≤ i ≤ N*_*p*_, 1 *≤j ≤N*_*s*_. Denote log_10_(cDNA equivalents/mL) viral load time series data for patients *i, j* via

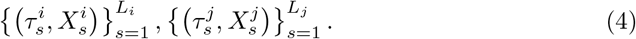

Our fitting procedure for an example pair of hosts is visually represented in Fig. 3.

1. For these hosts, fit the model to viral load data using the interior reflexive Newton-Rhapson method as implemented by MatLab’s `lsqcurvefit` function, minimizing

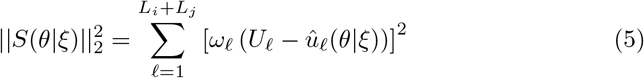

Where

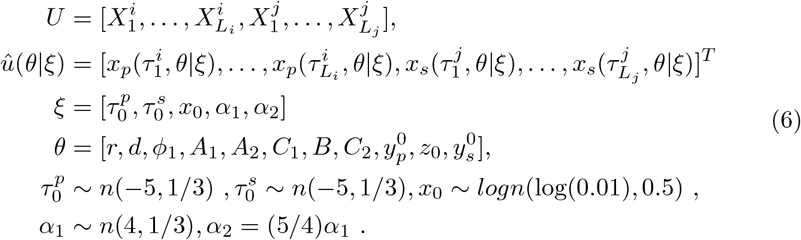
2. Here 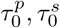 are the primary and secondary infection start times, respectively (based upon [3]). The distributions for *α*_1_ and *α*_2_, which represent the maximum replication enhancement and maximum clearance rates by the cross-reactive antibody, respectively, are calibrated so that reasonable immune response saturation levels occur, based upon [26], and so that viremia ends around the predicted timeframe [3, 27, 28].
3. For each pair of hosts, over 500 sets of these initialization parameters, we fit the model as described in step 1. We then retained the fit parameter values for which the residual norm was minimal subject to feasibility constraints max_*τ*_ *x*_*p*_(*τ*), max_*τ*_ *x*_*s*_(*τ*) *<* 20 log10(cDNA equivalents/mL) (chosen based upon examination of fits without these constraints). Over these iterations, each optimization is initialized by sampling 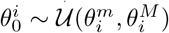, where 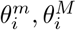 are the lower and upper bounds of the distributions specified in table 1.
4. To calculate viremia start, end, and duration times, we first need to define each of these in a way compatible with a mathematical model. Toward that end, we
5. characterize the start of viremia, or proliferation of the virus, as follows. *We say viremia has begun at time* 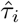*if (i)* 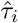 *falls in the interval from simulation start until the time at which x achieves its maximum (ii)* 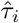 *is the first time at which x*(*τ*) *> E*_*i*_, where *E*_*i*_ is some appropriately chosen small positive value. Similarly, for viremia end *we say viremia has ended at time* 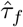*if (i)* 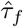*falls in the interval from peak viral load to the end of the simulation, (ii)* 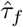*is the last time at which x*(*τ*) *> E*_*f*_, where *E*_*f*_ is some appropriately chosen small positive value. We have chosen *E*_*i*_ = *E*_*f*_ = 10 cDNA eq./mL for simplicity. We impose these conditions for each iteration of the simultaneous fitting procedure described above to find viremia start and end times and duration for primary and secondary DENV1, 2, and 3 patients.

**Fig 3.**
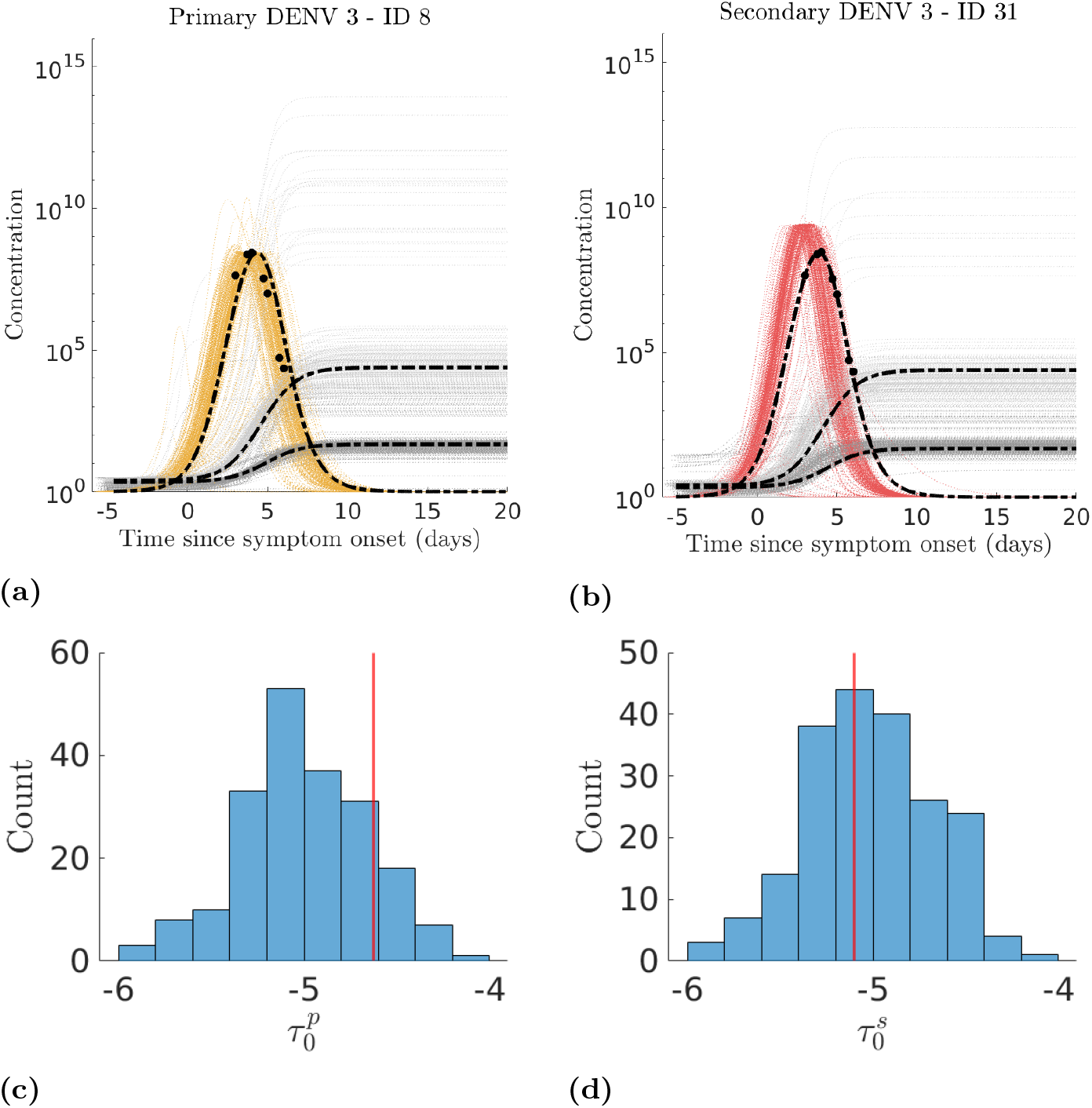
Visual representation of the simultaneous model fitting procedure for a single pair of primary and secondary hosts. **(a)** Example primary host data with fits. **(b)** Example secondary host data with fits. **(c)** Histogram of drawn mosquito bite times for the primary host. **(d)** Histogram of drawn mosquito bite times for the secondary host. The vertical red line represents the optimal infection start time, and bold dashed-dot lines represent the optimal fit among the biologically feasible fit parameter sets from the 500 sets of considered initialization parameters.

**Fig 4.**
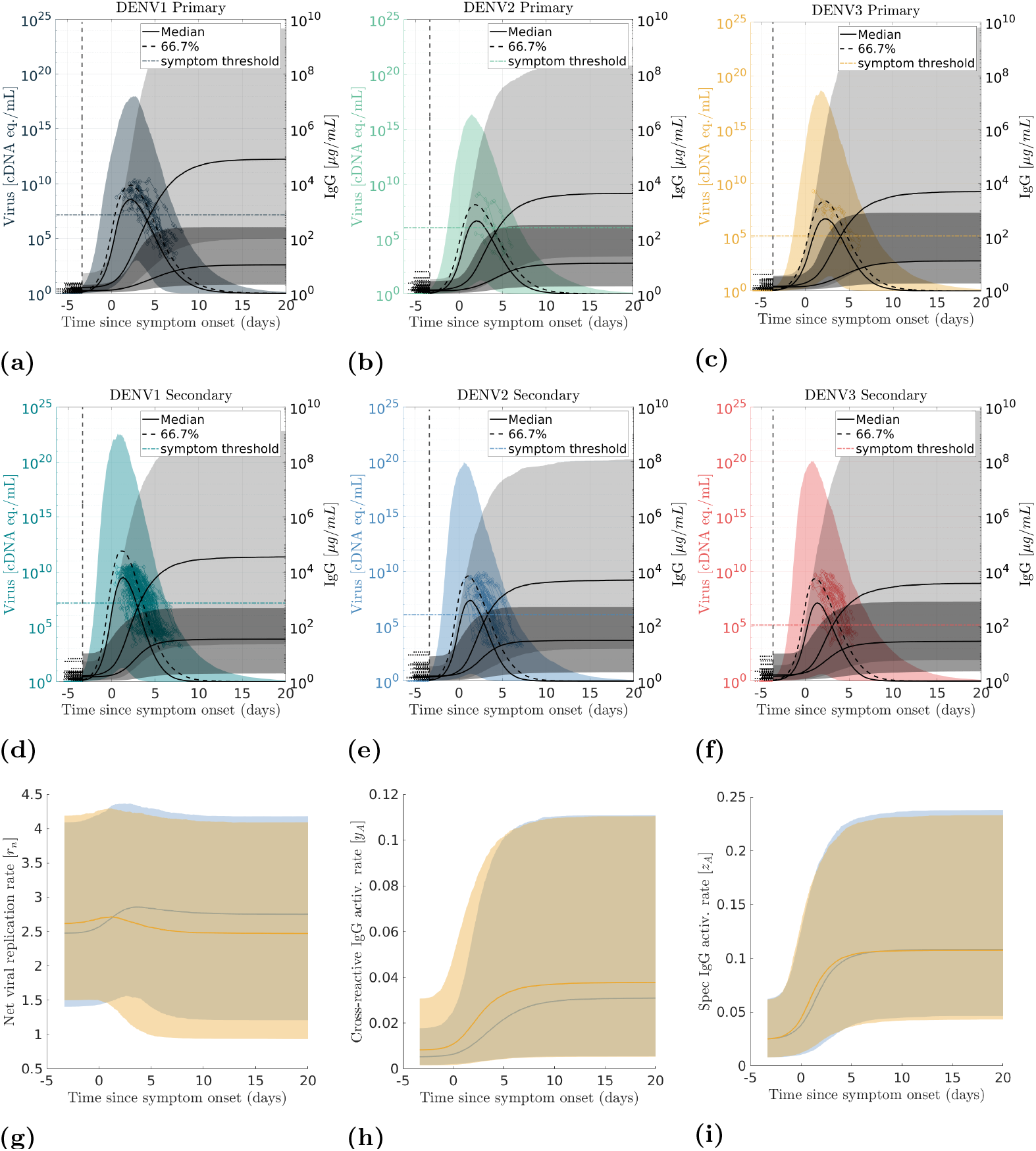
Model time trajectories and viral load time series. The model recapitulates the data well; the early spike in secondary viral load is due to Antibody-Dependent Enhancement via intermediate initial cross-reactive antibody level (see Fig. 6). **(a)-(c)** primary infection and **(d)-(f)** secondary infection. Colored points represent original virus time series data, and the shaded region represents 95 % confidence intervals. Dark grey represents cross-reactive antibody levels; light grey represents specific antibodies. Solid lines represent the median time trajectory for each model compartment. **(g)-(i)** Variation in activation/replication rates between primary and secondary infection. **(g)** net viral replication rate, **(h)** cross-reactive antibody activation rate.

**Table 1.**
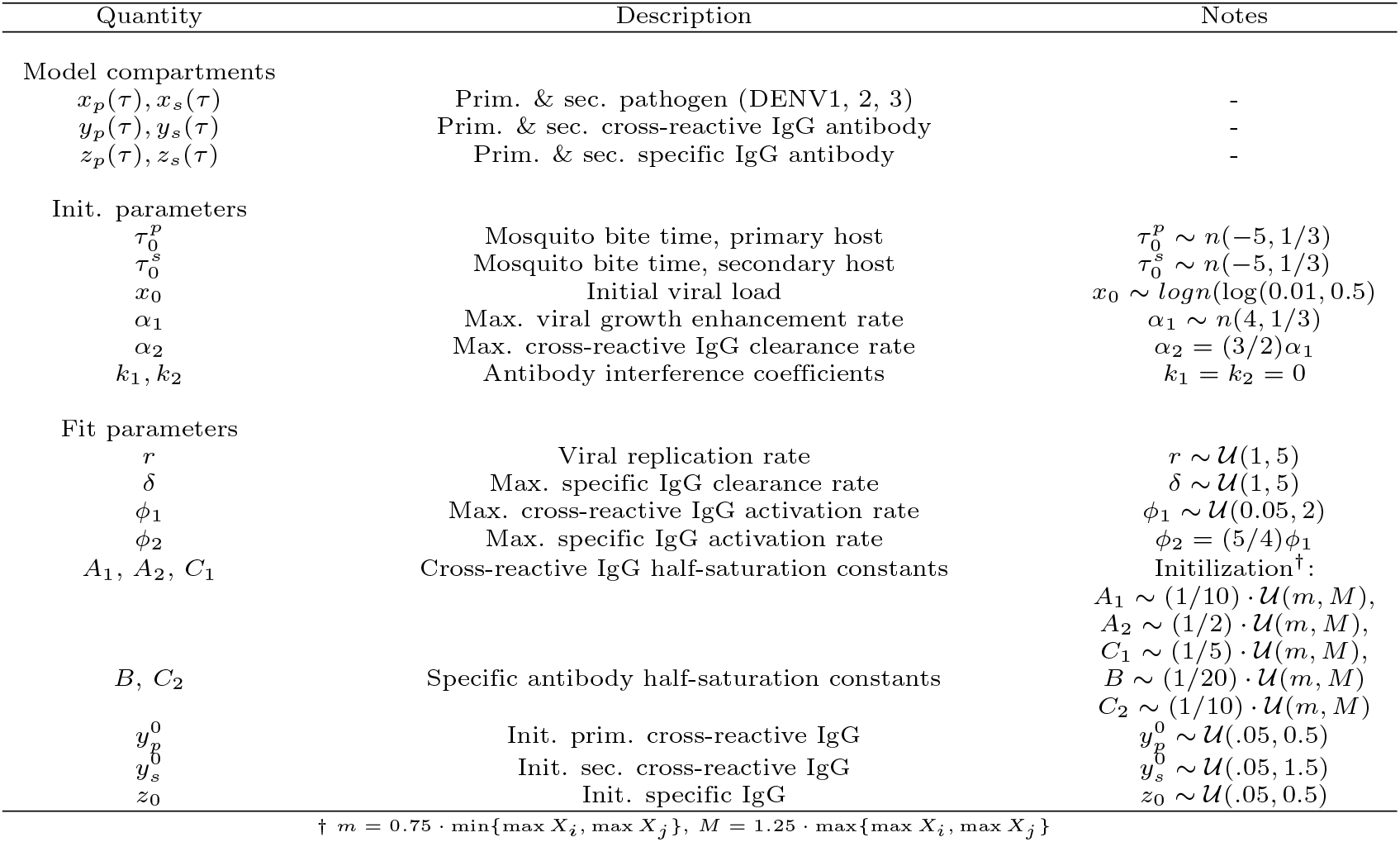
Key quantities for the model (3) with description. Due to the lack of associated antibody-kinetic data, some antibody parameter values and relationships are chosen based upon [3, 26]. Units for IgG antibodies are log_10_(*µg/mL*), and for viral load log_10_(*cDNA eq*.*/mL*).

### 3.4 Comparisons among serotype life history parameters

After fitting all possible combinations of hosts for each serotype as described above, we checked the life history parameters for significant differences among serotypes. To do this, we make use of Dunn’s multiple comparison tests with Holm’s correction for the p-values as implemented in the `scikit posthocs` package [32], which is a non-parametric pairwise posthoc test that crucially does not require that the size of the samples be equal. A low p-value is an indication that pairs of medians are significantly different. It is analogous to pairwise parametric tests but does not assume the underlying data distribution or the equality of variance between samples [33].

### 3.5 Simulation studies

Let *M* = log([Θ, ?]), where Θ, and Ξare the matrices whose rows are composed of the fit parameter estimates. Now assume that 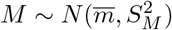, where, 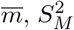 are the vector of sample column means and the sample covariance matrix of M (note that under this assumed distribution these are the maximum likelihood estimates for the distribution parameters of *M* [34]). Now, draw 10,000 parameter sets according to the above distribution subject to biological feasibility constraints max_*τ*_ *x*_*p*_, max_*τ*_ *x*_*s*_ *∈* (10^1^, 10^30^), max_*τ*_ *y*_*p*_, max_*τ*_ *y*_*s*_ *∈* (10^1*/*4^, 10^4^), sort the values, and remove the top and bottom 2.5% to obtain 95% confidence intervals for model parameters and resulting time trajectories.

This distribution was chosen to preserve the covariance structure of the fit model parameters, to ensure the sampled parameters’ positivity, and to allow for flexibility in the symmetry of the assumed distributions. Subsequently, we pair these results with data on long-term IgG kinetics to estimate the time frame of the intermediate risk window.

### 3.6 Estimating the intermediate risk window

To estimate the time frame of the intermediate risk window for elevated risk of severe secondary Dengue, which our model captures (Fig. 6), we consider a simple exponential decay model:

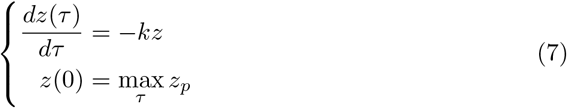

And then fit this model to each long-term post-primary infection IgG time series from [29]. We then assumed that

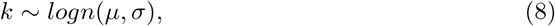

and obtained both unbiased point estimates and confidence intervals for *µ, σ*, using Matlab’s `lognfit` function. Next, over 10,000 replications, we uniformly randomly selected primary cross-reactive antibody saturation levels and drew a decay rate according to equation (8) (see Fig. 6). Finally, for each set of drawn values, we solved for the half-life, the entrance and exit times from the risk window, and the time spent in the risk window.

## 4. Results

### 4.1 Differences among viral life history parameters

Our model recapitulates the available data well (see Fig. 4), and our model fitting results indicate that there are significant differences in viral life history parameters among DENV1, 2, and 3 (see table 2, supplementary tables S1-S7 and supplementary figure S1).

**Table 2.** sample mean, median, and standard deviation for model parameters fit to data by serotype. See also supplementary tables S1-S7 and supplementary figure S1.

| Serotype | Statistic | $r$ | $\delta$ | $\phi_1$ | $A_1$ | $A_2$ | $C_1$ | $B$ | $C_2$ |
| --- | --- | --- | --- | --- | --- | --- | --- | --- | --- |
| DENV 1 | Mean | 2.32 | 4.06 | 0.11 | 1.06 | 4.94 | 2.24 | 0.44 | 0.85 |
|  | Median | 2.38 | 4.08 | 0.10 | 1.12 | 5.23 | 2.37 | 0.41 | 0.75 |
|  | Std. Dev. | 0.65 | 0.62 | 0.04 | 0.22 | 1.27 | 0.34 | 0.11 | 0.24 |
| DENV 2 | Mean | 1.99 | 3.94 | 0.12 | 0.84 | 4.43 | 1.86 | 0.36 | 0.70 |
|  | Median | 2.00 | 4.01 | 0.11 | 0.85 | 4.67 | 1.92 | 0.34 | 0.64 |
|  | Std. Dev. | 0.51 | 0.60 | 0.05 | 0.22 | 1.21 | 0.38 | 0.11 | 0.22 |
| DENV 3 | Mean | 2.03 | 3.87 | 0.11 | 0.91 | 4.22 | 2.03 | 0.36 | 0.70 |
|  | Median | 1.96 | 3.87 | 0.10 | 0.96 | 4.03 | 2.11 | 0.34 | 0.64 |
|  | Std. Dev. | 0.71 | 0.76 | 0.05 | 0.24 | 1.38 | 0.36 | 0.11 | 0.24 |
| Serotype | Statistic | $y_p^0$ | $z_0$ | $y_s^0$ | Prim bite | Sec bite | $x_0$ | $\alpha_1$ | |
| DENV 1 | Mean | 0.12 | 0.26 | 0.30 | -5.10 | -4.97 | 0.014 | 3.98 |  |
|  | Median | 0.09 | 0.27 | 0.12 | -5.15 | -4.96 | 0.011 | 3.98 |  |
|  | Std. Dev. | 0.08 | 0.13 | 0.37 | 0.48 | 0.49 | 0.009 | 0.36 |  |
| DENV 2 | Mean | 0.15 | 0.18 | 0.27 | -5.00 | -4.93 | 0.011 | 4.00 |  |
|  | Median | 0.07 | 0.16 | 0.11 | -4.98 | -4.92 | 0.009 | 4.00 |  |
|  | Std. Dev. | 0.13 | 0.12 | 0.31 | 0.45 | 0.48 | 0.007 | 0.36 |  |
| DENV 3 | Mean | 0.18 | 0.21 | 0.35 | -5.03 | -4.99 | 0.011 | 4.03 |  |
|  | Median | 0.12 | 0.20 | 0.21 | -5.05 | -5.02 | 0.009 | 4.04 |  |
|  | Std. Dev. | 0.14 | 0.11 | 0.34 | 0.39 | 0.35 | 0.007 | 0.32 |  |

Specifically, we note that DENV1 replicates more rapidly among serotypes than DENV2 or DENV3 [*r*]. A similar pattern holds for the half-saturation point of cross-reactive IgG clearance, half saturation of specific IgG clearance, and the half-saturation point of specific IgG activation [parameters *A*_2_, *B, C*_2_ respectively].

Though all serotypes had similar maximum cross-reactive activation rates, DENV1 reached half-saturation for replication enhancement at a relatively higher concentration than DENV3, and DENV2 achieved half-saturation at a lower concentration than either DENV1 or DENV3 [model parameter *A*_1_]. The same relative relationships hold for the half-saturation of cross-reactive IgG activation [parameter *C*_1_].

At the start of infection, DENV1 had both higher initial inoculum size and initial than either DENV2 or DENV3 and a larger amount of specific antibodies. In contrast, for both primary and secondary infections DENV1 and DENV2 have a lower concentration of cross-reactive antibodies than DENV3, and in secondary infections, DENV2 has a concentration of cross-reactive IgG that is intermediate between DENV1 and DENV3.

### 4.2 Differences between primary and secondary infection

We observe that the difference in initial IgG antibody levels results in differences between primary and secondary time and concentration-dependent activation rates (system of equations (2)). Specifically, we note that the median net viral replication rate, *r*_*n*_, and the specific antibody activation rate, *z*_*A*_, have switched in dominance during infection, with the former occurring around the onset of symptoms and just before peak viral load. This switch happens around symptom day five for *z*_*A*_, corresponding to the time around which the concentration of the IgG antibodies begins to increase rapidly (see Fig. 4).

In contrast, the median cross-reactive IgG activation rate, *y*_*A*_, is always greater for secondarily infected patients (Fig. 4g-i). As a direct consequence of these differences, our model recapitulates the clinically and empirically observed differences in the time course of events between primarily and secondarily infected DENV hosts.

Specifically, we note that the distributions for the start of viremia, time of maximum viral load, end of viremia, duration of viremia, and cumulative viral load all indicate similar upper bounds for their timeframe in both secondary and primary hosts but that secondary Dengue has both a lower mean and fatter left tail for each of these quantities (Fig. 5). Secondary patients tend to have a shorter duration of viremia and achieve peak viral load more rapidly than primary patients.

**Fig 5.**
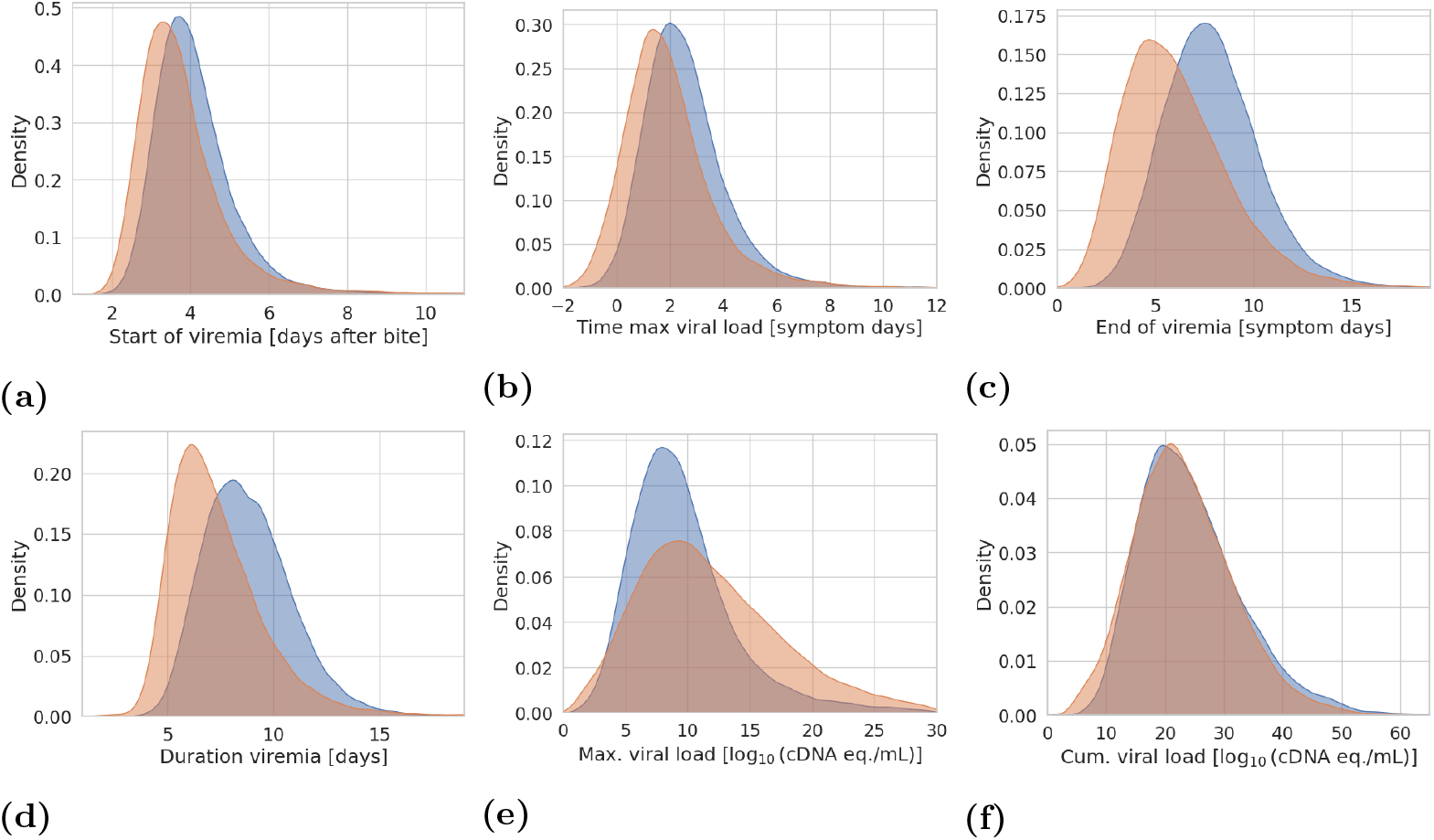
Differences in the time course of events between primary and secondary infection induced by differing initial cross-reactive antibody levels. Blue and orange curves are kernel density estimates from our primary and secondary infection simulation studies.

**Fig 6.**
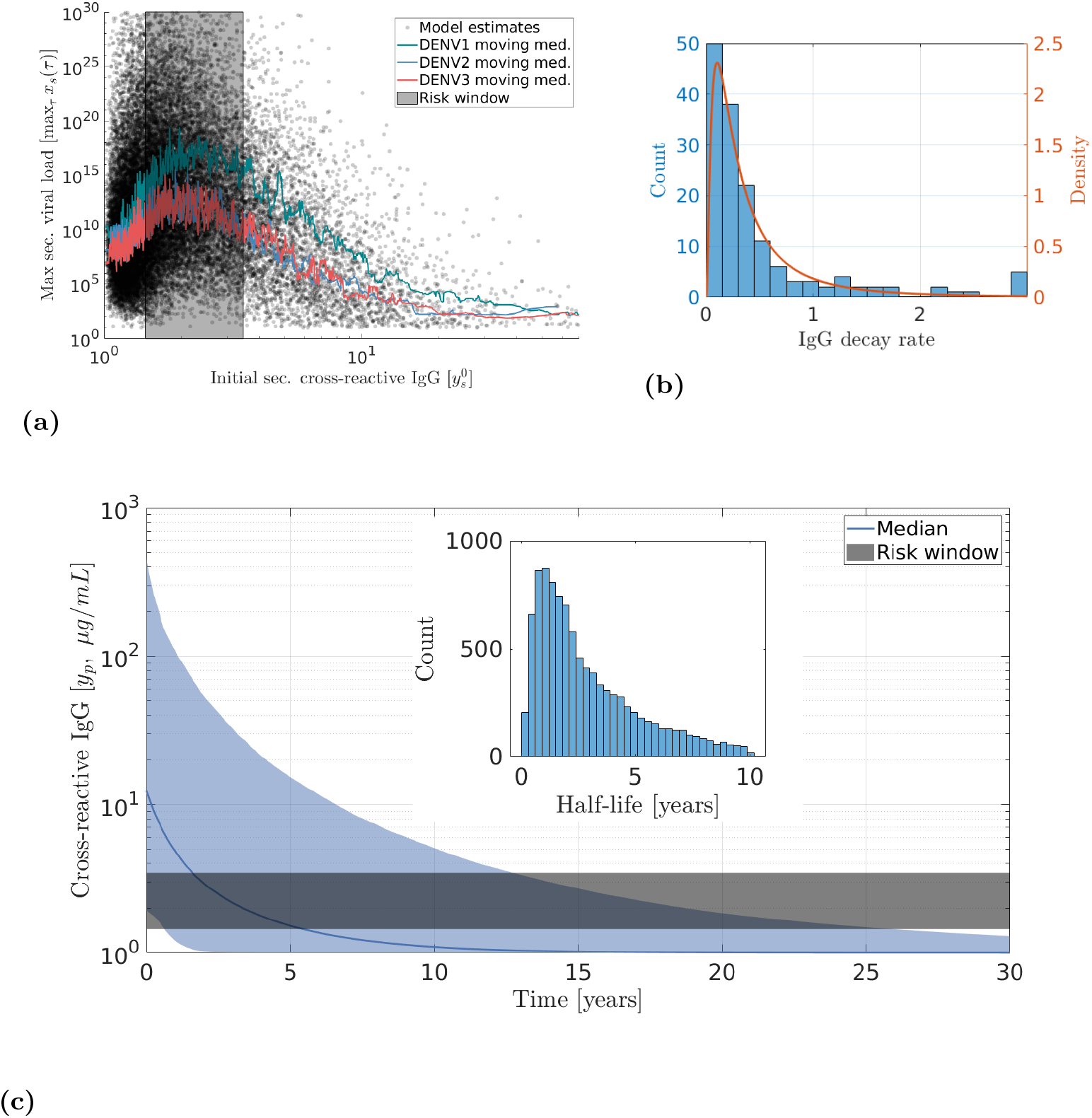
Intermediate risk window for elevated peak viral load in secondary dengue patients. **(a)** The relationship between initial cross-reactive antibody level and peak viral load from 10,000 generated parameter sets. The enhanced peak viral load region is approximately 1.45-3.47 *µ*g/mL. **(b)-(c)** Estimating the time frame of the intermediate risk window. **(b)** Decay rates estimated from long-term DENV antibody data **(c)** Decay trajectory of cross-reactive IgG antibodies **(insert)** IgG decay half-life distribution

In contrast to this for maximum viral load, which correlates to infection severity [2, 3], primary and secondary patients have similar mean viral loads, but the distribution of secondary maximum viral loads has fat tails, particularly to the right relative to primary maximum viral load. This difference reflects the reality that while secondarily infected hosts are at elevated risk for severe Dengue, it is not always the case that secondary infection is more severe than primary infection; indeed, in some circumstances, it may be less so [2, 3].

### 4.3 ADE effect and the intermediate risk window

Our simulation studies indicate that the cross-reactive antibody concentration associated with the intermediate risk window is consistently from approximately 1.45-3.47 *µ*g/mL across all three serotypes and that the chances of secondary infection occurring at all tail off significantly as concentrations exceed 10 *µg/mL* (see Fig. 6).

Significant variability exists in our estimates for the half-life of IgG antibodies (see supplementary table S8). This variability is likely due to variability among individual host immune responses, variation in primary infection severity, and the relatively infrequent collection of blood samples in the underlying data [29]. Nevertheless, we estimate a mean IgG half-life of 2.86 years and a median of 2.20 years (95% CI 0.0273-3.435 years). When combined with our estimates of primary cross-reactive IgG response saturation levels and the estimated range of concentrations associated with an elevated risk of severe Dengue, we estimate entrance into the intermediate risk window to occur a mean of 2.83 and a median of 1.67 years after primary infection (95% CI 0.0-12.5, see Fig. 6 and supplementary table S8). Once entered, we estimate that hosts spend a mean of 4.66 and a median of 3.58 years in this risk window (95% CI 0.42-14.08), ultimately exiting the risk window with a mean of 7.48 and a median of 5.42 years after primary infection (95% CI 0.67-25.42 years).

## 5 Discussion

From the perspective of data-driven modeling, there are multiple challenges to parameterizing a mechanistic model for DENV to sequential infections in humans. First, for various reasons, the beginning of infection and peak viral concentration are unobserved [3, 16, 27]. Second, due to the stochastic nature of secondary infections, primary and secondary *in vivo* patient data is generally unavailable from the same hosts [27]. Finally, immune and viral kinetic data are rarely simultaneously collected from patients.

We sought to mitigate these issues by simultaneously fitting the model (3) to all possible pairs of primary and secondary patients from the Hospital for Tropical Diseases (Ho Chi Minh City, Vietnam) between May 2007 and July 2008 (see Fig. 2, model (3), and section 3.2) while allowing for independent variation in infection start time among each host, and variation in initial viral load, and maximum cross-reactive response activation rate, selecting the optimal parameter set in terms of model residual for each paring from among 500 sets of parameter estimates given the randomly drawn infection start times. This fitting assumes that, on average, the only difference between a primary and secondary infection of the same serotype in distinct patients should be the difference in initial IgG antibody levels at the start of infection between the primarily and secondarily infected hosts, and the resulting concentration and time-dependent viral replication and IgG activation rates (see Fig. 4g-i). We found that our model recapitulated the data well (Fig. 4a-f) and captured clinically observed differences in the time course of events between primary and secondary DENV infections (Fig. 5).

Our results indicate significant differences in viral life history parameters among stereotypes (Figs. 5, S1). We note that DENV2 has a lower half-saturation point for both replication enhancement and cross-reactive IgG activation. These results may explain the empirical observation that secondary DENV2 is more likely to result in severe symptoms than the other serotypes (see [27] and supplementary table S9), at least for the data we considered.

Indeed, as we previously demonstrated [4], the range of initial cross-reactive IgG concentrations (from approximately 1.45-3.47 *µ*g/mL, see Fig. 6) where initial secondary cross-reactive antibody levels enhance viral replication is relatively small. Despite this, we find that for many primary-secondary pairs, the median secondary cross-reactive antibody level falls squarely within this range for our choice of other parameters (see tables 2, S1-S7). Notably, this effect is robust to the (co)variation allowed in the other model parameters as a part of our simulation studies (Fig. 6)a).

Finally, we utilized long-term NS1-specific IgG antibody data [29] to estimate both the half-lie of IgG antibodies and the time frame of the intermediate risk window relative to the onset of primary infection. We found that the half-life of IgG anti-bodes was between 0.33 and 8.62 years, with a mean of 2.86 years and a median of 2.20 years. Hosts entered the intermediate risk window between 0 and 12.5 years after primary infection (mean 2.83, median 1.67 years), stayed in this window between 0.42 and 14.08 years (mean 4.66 median 3.69), ultimately exiting the risk window between 0.67 and 25.42 years after primary infection (mean 7.48, median 5.42). These findings may at least partially explain the observed periodicity in the frequency of annually reported cases in countries where DENV is endemic (cf [35–37]), which are generally characterized by spikes in reported cases every two years, with occasional more significant peaks. Our results indicate that these patterns may be explained by cohorts of previously infected individuals entering into and out of the intermediate-risk window.

Given the relatively small sample size from a single time frame and in one country and other data limitations mentioned above, our findings are necessarily tentative and likely overestimate the prevalence of severe secondary Dengue, given that the data is collected from patients with sufficient systems to seek out treatment. Thus, we suggest several points that could generate the necessary data: (i) collection of both viral and immune kinetics from individual patients, (ii) when a patient is confirmed to have a secondary infection, query whether where/and when the primary infection occurred is known, (iii) record the severity of the infection (DF, DHF grades I, II). Such data would, perhaps, allow for both external validation of our results and a tighter estimate of the time frame of the intermediate risk window, allowing for optimally timed (re)vaccination campaigns and a corresponding reduction in the prevalence of severe Dengue.

## 6 Supporting information

**S1 Fig. Differences in median model parameters by serotype**. Daggers denote significant differences among the parameters estimated from the original data.

**S2 Fig. Sample correlation matrix for the pooled fit model parameters** from all combinations of primary and secondary hosts (*n* = 2700).

**S1 Table. Quantiles of the fit parameters for all possible DENV1 patient combinations**. (*N*_*p*_ *·N*_*s*_ = 2232)

**S2 Table. Quantiles of the fit parameters for all possible DENV2 patient combinations**. (*N*_*p*_ *·N*_*s*_ = 270)

**S3 Table. Quantiles of the fit parameters for all possible DENV3 patient combinations**. (*N*_*p*_ *·N*_*s*_ = 198)

**S4 Table. Dunn’s test for pairwise differences among DENV life history parameters with Holm’s correction for the p-values**

**S5 Table. Summary statistics for generated DENV1 parameter sets and resulting life history characteristics**

**S6 Table. Summary statistics for generated DENV2 parameter sets and resulting life history characteristics**

**S7 Table. Summary statistics for generated DENV3 parameter sets and resulting life history characteristics**

**S8 Table. Estimates for both IgG antibody half-life and the time frame of the intermediate risk window**.

**S9 Table. Prevalence of DHF among patients from [16, 27]**

## 7 Data availability statement

All generated parameter sets and code for associated analysis and figures are available on GitHub and deposited in citable format on the Zenodo repository [38]. The viral kinetics data is publicly available at [16]. The immune kinetics data is available upon request from the authors of [29].

## 8 Acknowledgments

The authors would like to thank Cameron Browne for his contributions to an earlier version of this manuscript and the authors of [29] for providing the long-term IgG data. A Zuckerman Foundation STEM leadership postdoctoral scholarship supports JCM. During the primary portion of this work, JCM and HG were partially supported by a US NSF RAPID grant (no. DMS-2028728) and NSF grant (no. DMS-1951759), and HG by a grant from the Simons Foundation/SFARI 638193.

