## Supplementary material for "Robust parameterization of a viral-immune kinetics model for sequential Dengue virus (DENV) infections with Antibody-Dependent Enhancement (ADE)": Supplmentary Materials

### Supplementary figures

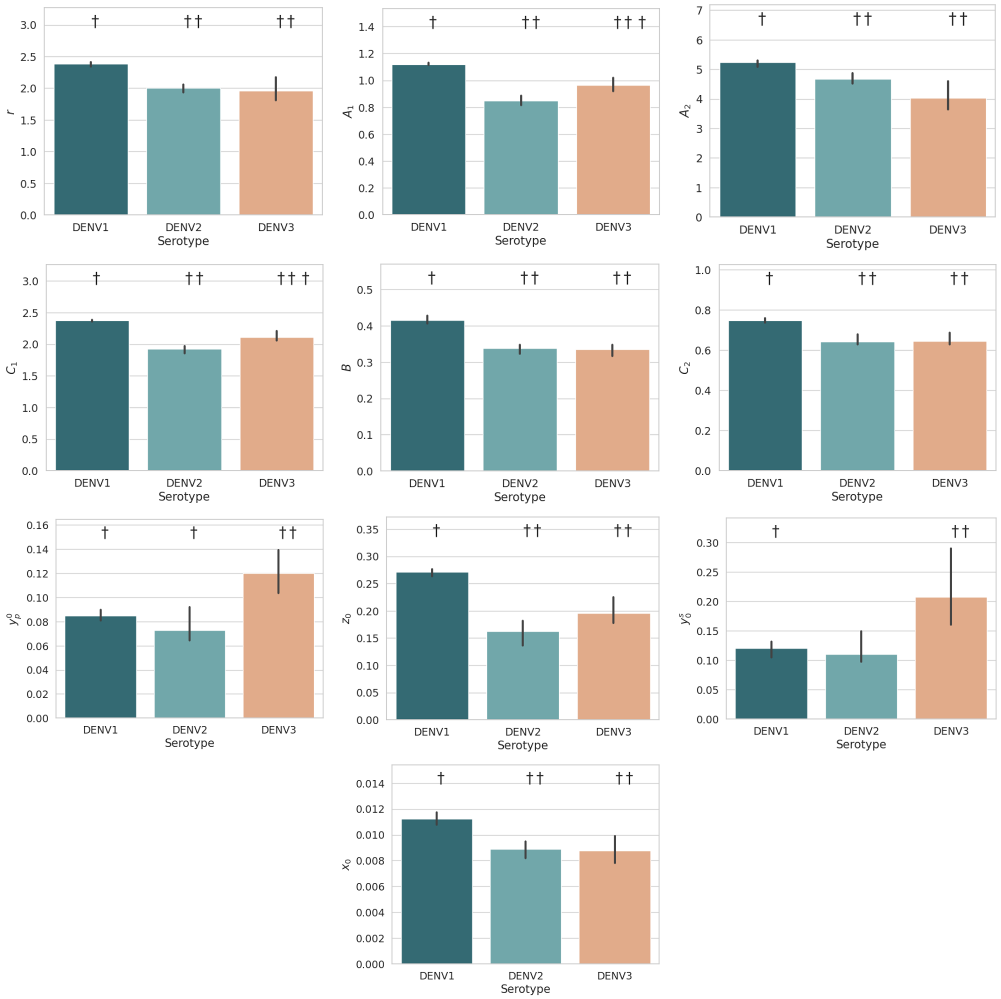


**Fig S1. Significant differences in median model parameters by serotype**. Daggers denote significant differences among the parameters estimated from the original data, p-value threshold $p < 0.01$. See also table S4.

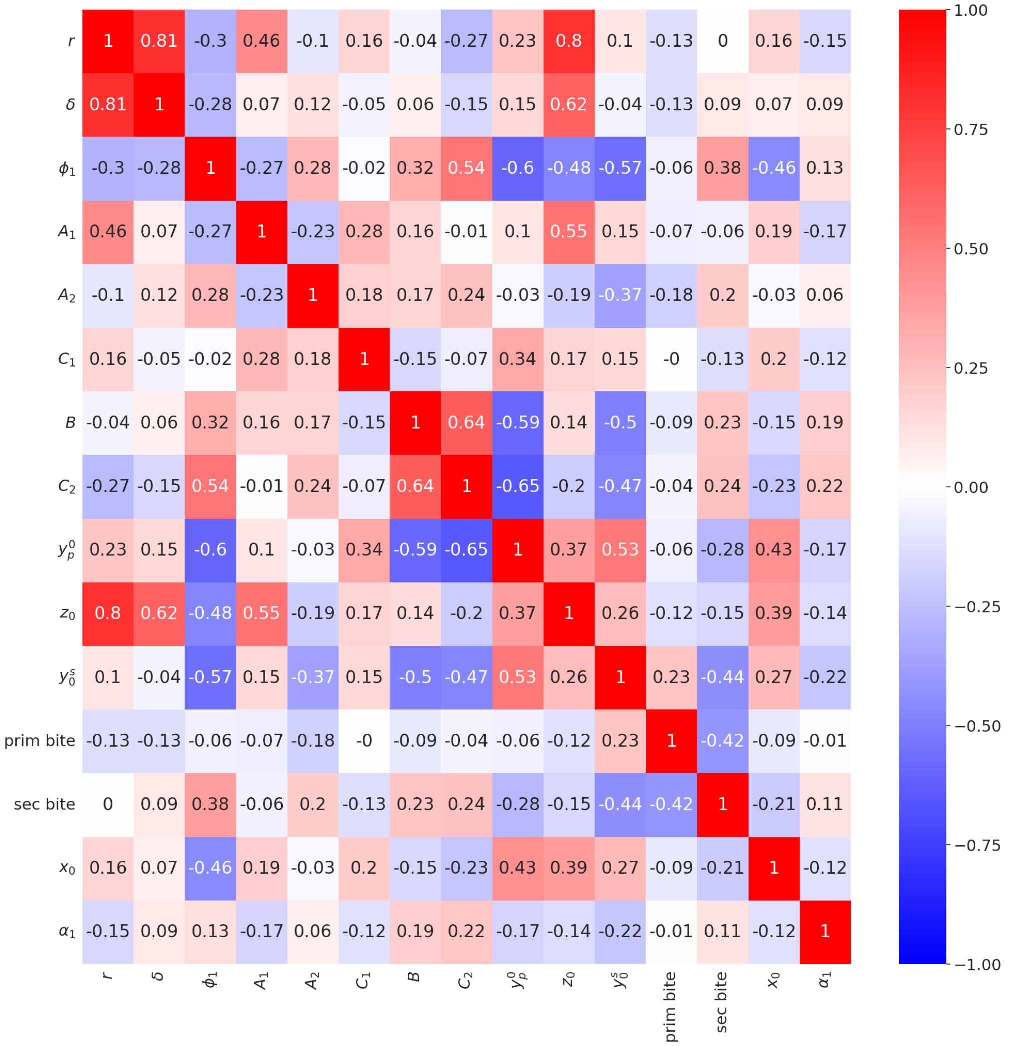


**Fig S2. Sample correlation matrix for the pooled fit model parameters** from all combinations of primary and secondary hosts ($n = 2700$).

### Supplementary tables

**Table S1. Quantiles of the fit parameters for all possible DENV1 patient combinations.** ($N_p \cdot N_s = 2232$)

| Parameter | 0.025 | 0.250 | 0.750 | 0.975 |
|---|---|---|---|---|
| $r$ | 1.156879 | 1.806015 | 2.816775 | 3.439883 |
| $\delta$ | 2.755604 | 3.622535 | 4.546330 | 4.999994 |
| $\phi_1$ | 0.050160 | 0.081903 | 0.129938 | 0.195097 |
| $A_1$ | 0.587112 | 0.901901 | 1.227669 | 1.301263 |
| $A_2$ | 2.774456 | 3.662303 | 6.138343 | 6.549419 |
| $C_1$ | 1.350786 | 2.059345 | 2.502100 | 2.623796 |
| $B$ | 0.274488 | 0.352784 | 0.536989 | 0.648435 |
| $C_2$ | 0.461271 | 0.667615 | 1.070051 | 1.282672 |
| $y_p^0$ | 0.050000 | 0.053721 | 0.177626 | 0.296872 |
| $z_0$ | 0.050000 | 0.144269 | 0.353048 | 0.499962 |
| $y_s^0$ | 0.050000 | 0.055474 | 0.292514 | 1.445492 |
| prim bite | -5.944389 | -5.466895 | -4.749057 | -4.139019 |
| sec bite | -5.889301 | -5.330456 | -4.587770 | -4.102201 |
| $x_0$ | 0.003382 | 0.007192 | 0.018431 | 0.037892 |
| $\alpha_1$ | 3.265393 | 3.728593 | 4.235263 | 4.646200 |

**Table S2.** Quantiles of the fit parameters for all possible DENV2 patient combinations ($N_p \cdot N_s = 270$)

| Parameter | 0.025 | 0.250 | 0.750 | 0.975 |
|---|---|---|---|---|
| $r$ | 1.153762 | 1.583247 | 2.332015 | 3.018820 |
| $\delta$ | 2.617172 | 3.568715 | 4.374961 | 4.936506 |
| $\phi_1$ | 0.051341 | 0.085787 | 0.140355 | 0.220874 |
| $A_1$ | 0.446996 | 0.672957 | 1.012819 | 1.192026 |
| $A_2$ | 2.068179 | 3.437962 | 5.519865 | 6.021909 |
| $C_1$ | 1.083018 | 1.634133 | 2.144545 | 2.388106 |
| $B$ | 0.177162 | 0.282088 | 0.418932 | 0.582597 |
| $C_2$ | 0.346085 | 0.534994 | 0.877465 | 1.145540 |
| $y_p^0$ | 0.050000 | 0.050311 | 0.222832 | 0.499987 |
| $z_0$ | 0.050000 | 0.067992 | 0.242868 | 0.434584 |
| $y_s^0$ | 0.050000 | 0.068977 | 0.385453 | 1.192750 |
| prim bite | -5.753507 | -5.324279 | -4.654235 | -4.150131 |
| sec bite | -5.795940 | -5.262608 | -4.606016 | -4.027669 |
| $x_0$ | 0.003012 | 0.005956 | 0.012519 | 0.032909 |
| $\alpha_1$ | 3.288006 | 3.735325 | 4.261630 | 4.654857 |

**Table S3.** Quantiles of the fit parameters for all possible DENV3 patient combinations ($N_p \cdot N_s = 198$)

| | 0.025 | 0.250 | 0.750 | 0.975 |
|---|---|---|---|---|
| $r$ | 1.000002 | 1.425017 | 2.500101 | 3.662658 |
| $\delta$ | 2.444188 | 3.342594 | 4.442169 | 4.999990 |
| $\phi_1$ | 0.050000 | 0.074797 | 0.136408 | 0.217399 |
| $A_1$ | 0.346119 | 0.725851 | 1.142207 | 1.211800 |
| $A_2$ | 1.730396 | 3.009632 | 5.664454 | 6.187551 |
| $C_1$ | 1.188218 | 1.807314 | 2.315267 | 2.466841 |
| $B$ | 0.173056 | 0.270756 | 0.440148 | 0.603205 |
| $C_2$ | 0.346074 | 0.509453 | 0.855807 | 1.209407 |
| $y_p^0$ | 0.050000 | 0.060496 | 0.249218 | 0.500000 |
| $z_0$ | 0.050001 | 0.116135 | 0.278276 | 0.443455 |
| $y_s^0$ | 0.050000 | 0.086506 | 0.553623 | 1.355854 |
| prim bite | -5.748548 | -5.334385 | -4.750386 | -4.321240 |
| sec bite | -5.666117 | -5.227822 | -4.743739 | -4.344739 |
| $x_0$ | 0.003857 | 0.005704 | 0.013499 | 0.028534 |
| $\alpha_1$ | 3.450789 | 3.783422 | 4.284942 | 4.660997 |

**Table S4. Dunn's test for pairwise differences among DENV life history parameters with Holm's correction for the p-values.**

| | DENV1 vs DENV2 | DENV1 vs DENV3 | DENV2 vs DENV3 |
|---|---|---|---|
| $r$ | 0.000 | 0.000 | 0.349 |
| $\delta$ | 0.012 | 0.005 | 0.535 |
| $\phi_1$ | 0.082 | 0.855 | 0.339 |
| $A_1$ | 0.000 | 0.000 | 0.001 |
| $A_2$ | 0.000 | 0.000 | 0.188 |
| $C_1$ | 0.000 | 0.000 | 0.000 |
| $B$ | 0.000 | 0.000 | 0.933 |
| $C_2$ | 0.000 | 0.000 | 0.747 |
| $y_p^0$ | 0.965 | 0.000 | 0.002 |
| $z_0$ | 0.000 | 0.000 | 0.024 |
| $y_s^0$ | 0.051 | 0.000 | 0.013 |
| Prim bite | 0.001 | 0.033 | 0.606 |
| Sec bite | 0.652 | 0.652 | 0.493 |
| $x_0$ | 0.000 | 0.000 | 0.970 |
| $\alpha_1$ | 0.686 | 0.159 | 0.686 |

**Table S5. Summary statistics for generated DENV1 parameter sets and resulting life history characteristics**

| Quantity | Mean | Median | Std. Dev | 95% CI |
|---|---|---|---|---|
| $r$ | 2.291979 | 2.193582 | 0.676051 | (1.224, 3.847) |
| $\delta$ | 4.006229 | 3.952808 | 0.634418 | (2.909, 5.354) |
| $\phi_1$ | 0.108061 | 0.101750 | 0.036645 | (0.056, 0.197) |
| $A_1$ | 1.062751 | 1.037723 | 0.238478 | (0.663, 1.599) |
| $A_2$ | 4.838512 | 4.656475 | 1.327125 | (2.764, 7.937) |
| $C_1$ | 2.239811 | 2.204488 | 0.397613 | (1.563, 3.118) |
| $B$ | 0.437257 | 0.422038 | 0.117321 | (0.255, 0.706) |
| $C_2$ | 0.845811 | 0.811304 | 0.246822 | (0.468, 1.417) |
| $y_p^0$ | 0.119353 | 0.100327 | 0.076913 | (0.03, 0.314) |
| $z_0$ | 0.257952 | 0.212000 | 0.178264 | (0.058, 0.722) |
| $y_s^0$ | 0.266736 | 0.174503 | 0.262985 | (0.03, 1.015) |
| prim bite | -5.063023 | -5.045285 | 0.467077 | (-6.028, -4.205) |
| sec bite | -4.996407 | -4.976413 | 0.468843 | (-5.982, -4.142) |
| $x_0$ | 0.013854 | 0.011245 | 0.009947 | (0.003, 0.039) |
| $\alpha_1$ | 3.963107 | 3.946888 | 0.360848 | (3.298, 4.705) |
| max prim | 10.376478 | 9.684185 | 3.824948 | (4.922, 21.194) |
| t. max prim | 2.410032 | 2.222426 | 1.415578 | (0.259, 5.687) |
| max sec | 13.357556 | 12.618175 | 5.754180 | (4.226, 26.394) |
| t. max sec | 1.694722 | 1.587042 | 1.415449 | (-0.682, 4.859) |
| cum prim | 29.268799 | 28.257210 | 8.451618 | (15.571, 48.698) |
| cum sec | 27.063531 | 26.733110 | 8.292870 | (11.538, 44.858) |
| v. start prim | 3.900457 | 3.795516 | 0.828147 | (2.669, 5.754) |
| v. start sec | 3.478321 | 3.349915 | 0.877232 | (2.273, 5.547) |
| v end prim | 8.801382 | 8.662145 | 2.175654 | (4.931, 13.587) |
| v end sec | 6.340252 | 5.987042 | 2.873111 | (1.962, 12.787) |
| dur v. prim | 9.963948 | 9.800000 | 1.732100 | (6.9, 13.8) |
| dur v. sec | 7.858339 | 7.500000 | 2.357930 | (4.6, 13.4) |
| p. cross IgG sat | 1.119431 | 1.043276 | 0.557644 | (0.321, 2.418) |

**Table S6. Summary statistics for generated DENV2 parameter sets and resulting life history characteristics**

| Quantity | Mean | Median | Std. Dev | 95% CI |
|---|---|---|---|---|
| $r$ | 1.976850 | 1.913229 | 0.523171 | (1.139, 3.162) |
| $\delta$ | 3.866614 | 3.822855 | 0.609707 | (2.806, 5.193) |
| $\phi_1$ | 0.119828 | 0.110666 | 0.048235 | (0.053, 0.238) |
| $A_1$ | 0.851099 | 0.820450 | 0.237395 | (0.472, 1.399) |
| $A_2$ | 4.332854 | 4.148872 | 1.353291 | (2.279, 7.544) |
| $C_1$ | 1.820569 | 1.773162 | 0.426030 | (1.121, 2.783) |
| $B$ | 0.353889 | 0.337892 | 0.111533 | (0.186, 0.615) |
| $C_2$ | 0.697531 | 0.656042 | 0.240582 | (0.35, 1.286) |
| $y_p^0$ | 0.149664 | 0.118098 | 0.116076 | (0.027, 0.462) |
| $z_0$ | 0.182379 | 0.147809 | 0.131331 | (0.038, 0.532) |
| $y_s^0$ | 0.255235 | 0.180794 | 0.227510 | (0.031, 0.894) |
| prim bite | -4.999168 | -4.979418 | 0.450153 | (-5.944, -4.18) |
| sec bite | -4.924221 | -4.895281 | 0.489748 | (-5.936, -4.037) |
| $x_0$ | 0.010516 | 0.008788 | 0.006916 | (0.003, 0.029) |
| $\alpha_1$ | 3.987348 | 3.975356 | 0.357689 | (3.338, 4.724) |
| max prim | 8.954830 | 8.198531 | 4.013620 | (3.278, 19.098) |
| t. max prim | 2.478134 | 2.233290 | 1.629991 | (0.031, 6.325) |
| max sec | 11.001750 | 10.260930 | 5.257983 | (2.842, 23.395) |
| t. max sec | 1.873664 | 1.624789 | 1.736987 | (-0.694, 6.006) |
| cum prim | 21.524284 | 20.310545 | 7.450893 | (10.509, 39.65) |
| cum sec | 21.311819 | 20.475310 | 7.727192 | (8.413, 38.732) |
| v. start prim | 4.134943 | 3.957711 | 1.125426 | (2.596, 6.836) |
| v. start sec | 3.738519 | 3.520085 | 1.148157 | (2.289, 6.623) |
| v end prim | 7.399864 | 7.123659 | 2.516453 | (3.411, 13.286) |
| v end sec | 5.907204 | 5.465397 | 2.786907 | (1.824, 12.525) |
| dur v. prim | 8.264089 | 8.000000 | 2.062591 | (5.2, 13.1) |
| dur v. sec | 7.092906 | 6.700000 | 2.182534 | (4.1, 12.3) |
| p. cross IgG sat | 1.167961 | 1.120538 | 0.609087 | (0.279, 2.514) |

**Table S7. Summary statistics for generated DENV3 parameter sets and resulting life history characteristics**

| Quantity | Mean | Median | Std. Dev | 95% CI |
|---|---|---|---|---|
| $r$ | 1.981841 | 1.878843 | 0.705977 | (0.921, 3.608) |
| $\delta$ | 3.760887 | 3.684025 | 0.809708 | (2.418, 5.579) |
| $phi_1$ | 0.115634 | 0.105807 | 0.048546 | (0.05, 0.24) |
| $A_1$ | 0.934020 | 0.891967 | 0.298669 | (0.48, 1.647) |
| $A_2$ | 4.193670 | 3.957687 | 1.481265 | (2.04, 7.757) |
| $C_1$ | 2.007198 | 1.960044 | 0.421257 | (1.308, 2.934) |
| $B$ | 0.354672 | 0.335958 | 0.117103 | (0.181, 0.633) |
| $C_2$ | 0.707404 | 0.662131 | 0.258177 | (0.342, 1.339) |
| $y_p^0$ | 0.177787 | 0.140939 | 0.130981 | (0.035, 0.534) |
| $z_0$ | 0.207795 | 0.173164 | 0.137973 | (0.048, 0.569) |
| $y_s^0$ | 0.307528 | 0.224611 | 0.259344 | (0.038, 0.997) |
| prim bite | -5.038788 | -5.025645 | 0.394930 | (-5.863, -4.295) |
| sec bite | -4.981854 | -4.969269 | 0.350944 | (-5.708, -4.337) |
| $x_0$ | 0.010531 | 0.009035 | 0.006396 | (0.003, 0.027) |
| $\alpha_1$ | 4.023366 | 4.009277 | 0.320940 | (3.425, 4.682) |
| max prim | 9.470478 | 8.351752 | 4.833696 | (3.21, 22.686) |
| t. max prim | 2.858755 | 2.632505 | 1.639337 | (0.389, 6.61) |
| max sec | 10.913680 | 10.023515 | 5.586838 | (2.418, 24.335) |
| t. max sec | 2.323375 | 1.909782 | 2.068765 | (-0.391, 7.41) |
| cum prim | 22.254008 | 20.888660 | 8.243816 | (10.093, 42.498) |
| cum sec | 21.212673 | 20.410500 | 8.117221 | (7.254, 40.144) |
| v. start prim | 4.322460 | 4.167334 | 1.131322 | (2.694, 6.896) |
| v. start sec | 3.989772 | 3.722126 | 1.294778 | (2.407, 7.31) |
| v end prim | 7.580824 | 7.273663 | 2.544264 | (3.609, 13.563) |
| v end sec | 6.192895 | 5.688572 | 2.859302 | (2.263, 13.374) |
| dur v. prim | 8.297153 | 7.900000 | 2.137133 | (5.3, 13.502) |
| dur v. sec | 7.184976 | 6.700000 | 2.233580 | (4.3, 12.9) |
| p. cross IgG sat | 1.220473 | 1.117577 | 0.724751 | (0.277, 2.91) |

**Table S8. Estimates for both IgG antibody half-life and the time frame of the intermediate risk window (in years).**

| Quantity | Mean | Median | Std. Dev. | 95$ CI |
|---|---|---|---|---|
| Entrance | 2.83 | 1.67 | 3.39 | (0, 12.50) |
| Exit | 7.48 | 5.42 | 6.49 | (0.67, 25.42) |
| Stay | 4.66 | 3.58 | 3.69 | (0.42, 14.08) |
| Half-life | 2.86 | 2.20 | 2.21 | (0.33, 8.62) |

**Table S9. Prevalence of DHF among patients from [1, 2]**

| Infection | DENV1 | DENV2 | DENV3 |
|---|---|---|---|
| Primary | 16.7% (3/18) | 16.7% (1/6) | 0% (0/6) |
| Secondary | 26.6% (33/124) | 46.7% (21/45) | 30.3% (10/33) |
